# Physical activity regulates the immune response to breast cancer by a hematopoietic stem cell-autonomous mechanism

**DOI:** 10.1101/2023.09.30.560299

**Authors:** Lyne Khair, Katherine Hayes, Amanda Tutto, Amruta Samant, Lindsay Ferreira, Tammy T. Nguyen, Michael Brehm, Louis M. Messina

## Abstract

Physical activity is a modifiable lifestyle factor that is associated with a decreased risk for the development of breast cancer. While the exact mechanisms for the reduction in cancer risk due to physical activity are largely unknown, it is postulated that the biological reduction in cancer risk is driven by improvements in inflammation and immune function with exercise. Hematopoietic stem cells (HSCs) are the progenitor for all of the cells of the immune system and are involved in cancer immunosurveillance through differentiation into cytotoxic cell population. In this study, we investigate the role of physical activity (PA) in a spontaneously occurring model of breast cancer over time, with a focus on tumor incidence, circulating and tumor-infiltrating immune cells as well gene expression profiles of tumors and hematopoietic stem cells. Furthermore, we show that, in addition to a direct effect of PA on the immune cells of tumor-bearing mice, PA reduces the oxidative stress in HSCs of wildtype and tumor-bearing mice, and by doing so, alters the differentiation of the HSCs towards T cells in order to enhance cancer immunosurveillance.

## INTRODUCTION

Cancer is the second leading cause of death in the United States behind only heart disease^23^. Breast cancer is the most common cancer in women and is responsible for the most cancer deaths in women worldwide. Among the different subtypes of breast cancers, triple-negative breast cancer (TNBC) is more likely to affect younger women, who are African American or Latina, and those with a BRCA1 gene mutation^10, 21, 43, 46^. Triple-negative breast cancer has fewer treatment options than other types of invasive breast cancer. It is characterized by lack of expression of the estrogen receptor (ER), progesterone receptor (PR) and absence of *ERBB2* (also known as *HER2*) amplification, which makes hormone therapy or targeted drugs less likely to work. While surgery and chemotherapy are treatments of choice, radiation is sometimes prescribed as an option depending on certain features of the tumor^1^, as well as PARP inhibitors and checkpoint inhibitors^28^.

Physical activity is a modifiable lifestyle factor that is associated with a decreased risk for the development of cancer^35, 49^. However, the mechanisms for the protective effects of physical activity against cancer development and progression remain unknown. It is believed that physical activity improves immune cell function to enhance the cancer-fighting ability of the immune system. Reductions in cancer risk with physical activity may vary depending on the type of cancer. An analysis that examined 1.44 million people from 12 prospective study cohorts showed that people who were in the top 90% for leisure time physical activity had a risk reduction for 13 out of the 26 cancers examined, including breast, lung, colon, liver, endometrial, myeloid leukemia, myeloma, and head and neck cancers^35^. While the risk for many types of cancer may be reduced with regular participation in physical activity, the strongest evidence for cancer risk reduction with physical activity has been shown in breast and colon cancers^5^.

There is strong epidemiological evidence for the relationship between physical activity and a decreased risk of breast cancer. Higher levels of total physical activity, including recreational activities, deliberate exercise, and non-recreational physical activity, has been shown to confer a 25% reduction in risk of breast cancer relative to the lowest levels of physical activity^5^. A systematic review and meta-analysis examined the effect of recreational physical activity on breast cancer risk stratified by menopausal status^37^. Pizot et al.^42^ also reported that increased levels of physical activity reduce the risk of cancer, regardless of menopausal status and tumor hormone receptor status^42^. Overall, there is robust evidence showing the positive effects of exercise on reducing the risk of breast cancer.

While the exact mechanisms for the reduction in cancer risk due to physical activity are largely unknown, it is postulated that the biological reduction in cancer risk is driven by improvements in inflammation and immune function with exercise^19, 24^. A reduction in adiposity concomitant with regular physical activity lowers the levels of systemic inflammation in people that are chronically active^14, 32^. Furthermore, physical activity may protect against DNA oxidative damage, promote DNA repair, and stimulate skeletal muscle myokine release^32^.

Acute and chronic physical activity has a vast range of physiological effects on immune cells, influencing both cell number and cell function. In general, moderate physical activity has been associated with increased T cell proliferation^38, 45^, decreased levels of senescent T cells^50^, lower levels of inflammatory cytokines^40^, and greater NK cell cytotoxicity^3, 38, 41^. Increases in circulating cell numbers and cell cytotoxicity may be important in immune cell surveillance against cancer. Single bouts of physical activity can directly increase the quantity and function of cytotoxic cells, but it is the cumulative effect of regular physical activity that has been associated with a reduced risk for cancer in humans. Therefore, it is important to understand the role of chronic exercise on the cancer immunosurveillance function on immune cells. Most studies that have evaluated the effects of chronic exercise on immune system function have examined athletes to assess the potential immunosuppressive effects of intense exercise^57^, or they have studied elderly populations to better understand ways to preserve immunity throughout the life span^47^. Few studies have investigated the effects of moderately intense physical activity on immune cell function, with even fewer studies examining the effect of moderate intensity exercise training on cancer immunosurveillance.

Hematopoietic stem cells (HSCs) are the progenitor for all of the cells of the immune system; hematopoietic stem cells are primitive cells found primarily in the bone marrow, and to a lesser extent, in extramedullary sites including the spleen and thymus. HSCs are characterized by self-renewal as well as differentiation into myeloid and lymphoid lineages. Since HSCs are the progenitor for all of the cells of the immune system, it stands to reason that HSCs are involved in cancer immunosurveillance through differentiation into cytotoxic cell population. However, to date, only one study has demonstrated a role for HSC differentiation in the initiation and progression of cancer^55^. Tie et al recently demonstrated a role for HSC lineage priming in cancer immunosurveillance. In that study, hypercholesterolemia was found to induce oxidant stress-dependent epigenetic changes in HSCs that skewed their differentiation away from cancer surveying NKT and γδ T cells, resulting in increased incidence and histological severity of colon cancer in mice^55^. To our knowledge, this is the only study demonstrating that oxidant stress in HSCs can affect cancer immunosurveillance. However, a generalized role for oxidative stress in cancer development is more firmly established.

In this study, we investigate the role of physical activity (PA) in a spontaneously occurring model of breast cancer over time, with a focus on tumor incidence, circulating and tumor-infiltrating immune cells as well gene expression profiles of tumors and hematopoietic stem cells. Furthermore, we show that, in addition to a direct effect of PA on the immune cells of tumor-bearing mice, PA reduces the oxidative stress in HSCs of wildtype and tumor-bearing mice, and by doing so, alters the differentiation of the HSCs toward CD4+ T cells in order to enhance cancer immunosurveillance.

## RESULTS

### Physical activity decreases oxidant stress in hematopoietic stem cells and leads to increased circulating neutrophils in wildtype mice

To test the role of physical activity on oxidant stress levels in murine hematopoietic stem cells, we placed C57BL6/J female wildtype (WT) mice at 3 weeks of age in cages fitted with running wheels. The physically active (PA) mice were monitored, and daily running distances were recorded (Figure 1A). Sedentary (SED) age-matched mice were placed in regular cages. Following 12 weeks of physical activity, mice were sacrificed, and their bone marrow was isolated and enriched for lineage negative cells. Hematopoietic stem cells (HSCs) were defined as Lineage-c-kit+ sca1+ (LSK) and were purified by fluorescence-activated cell sorting (FACS). The resulting cell populations were analyzed for oxidant stress levels, as measured by mean fluorescence intensity using a DCF (Dichlorofluorescin-diacetate) assay to measure reactive oxygen species (Abcam). The results of this experiment showed a significant decrease in reactive oxygen species (ROS) in HSCs derived from PA mice (p=0.026) (Figure 1B, C).

**Figure 1.**
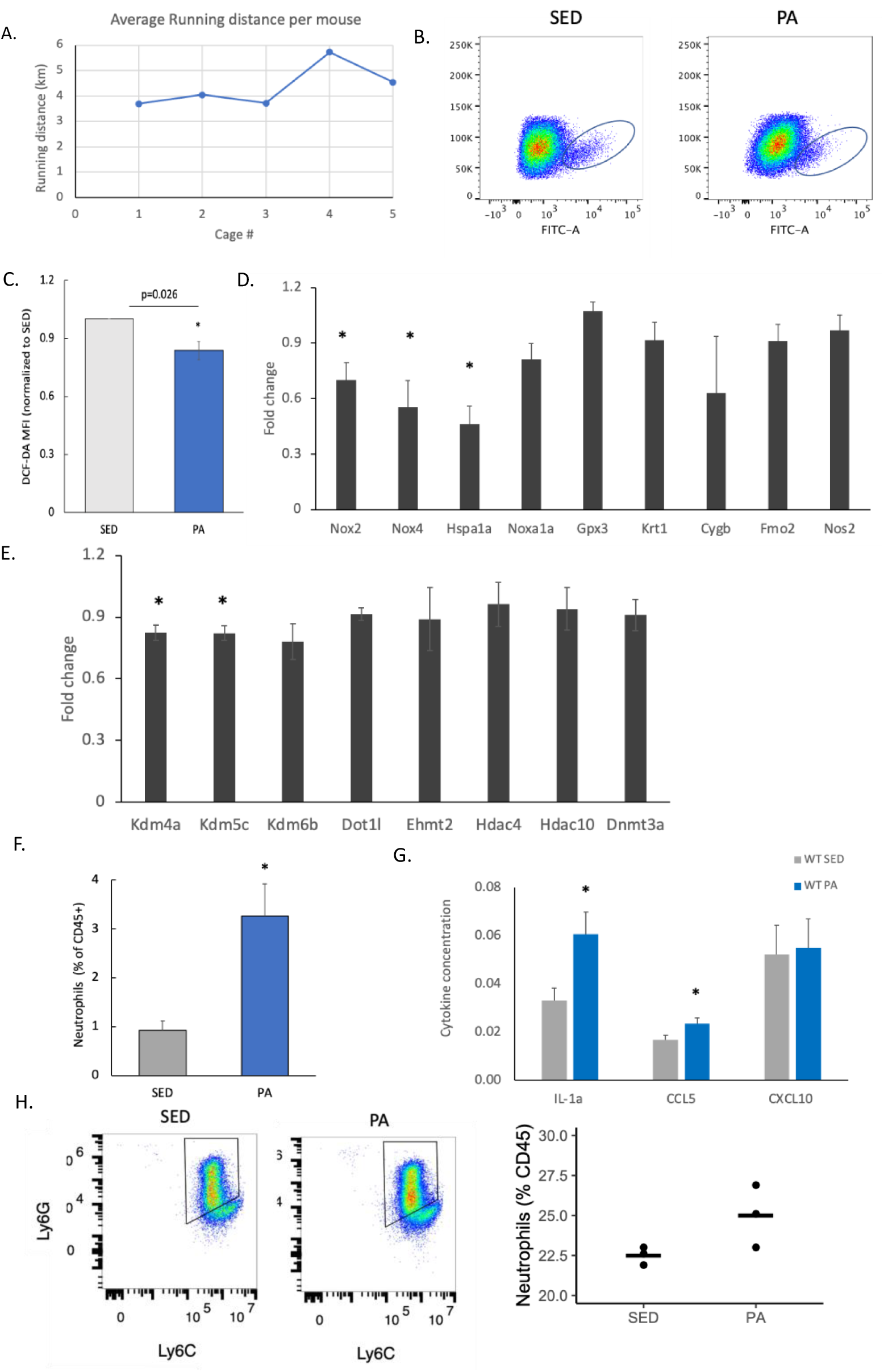
Physical activity decreases oxidant stress in hematopoietic stem cells and leads to increased circulating neutrophils in wildtype mice. **A.** Average running distance per mouse (n=5). **B.** Representative fluorescence analysis of DCF-DA assay by flow cytometry in HSCs from WT SED and PA mice. **C.** Quantification of the DCF-DA mean fluorescence intensity (MFI) in HSCs from WT SED and PA mice (n=5). **D.** Gene expression quantification by qPCR of oxidative stress enzymes in PA HSCs, normalize to SED HSC levels (n=4). **E.** Gene expression quantification by qPCR of epigenetic enzymes in PA HSCs, normalize to SED HSC levels (n=4). **F.** Quantification of circulating neutrophils in WT SED and PA mice, as a % of CD45+ cells (n=5). **G.** Cytokine concentration in serum from WT SED and PA mice (n=6-9). **H.** Representative flow cytometry analysis of neutrophils (CD45+ Ly6C+ Ly6G+) derived from in vitro differentiation of HSCs from WT SED and PA mice. I. Quantification of neutrophils derived from in vitro differentiation of HSCs from WT SED and PA mice (n=3). (Error bars=SEM; *, p,0.05; 2-tailed Student’s t-test).

In order to identify the enzymes responsible for the ROS decrease in HSCs from PA mice, we used the RT² Profiler™ PCR Array Mouse Oxidative Stress and Antioxidant Defense kit (Qiagen). We subsequently confirmed the top differentially expressed genes using qPCR. In 4 independent experiments, as shown in Figure 1D, the expression levels of Nox2, Nox4 and Hspa1a were all significantly decreased in HSCs from PA mice (p=0.04, 0.05 and 0.03, respectively). Taken together, these results show that physical activity decreases oxidative stress in HSCs, a previously unreported finding.

In our previous studies, we found that changes in oxidant stress enzymes lead to epigenetic imprinting in HSCs. To test whether epigenetic modifying enzymes were dysregulated in HSCs following physical activity, we used the RT² Profiler™ PCR Array Mouse Epigenetic Chromatin Modification Enzymes kit and confirmed the expression levels of the top differentially expressed genes using qPCR. We found that the expression levels of Kdm4a and Kdm5c were significantly decreased in HSCs (p=0.01) (Figure 1E). These results indicate a potential role for physical activity in the epigenetic imprinting of HSCs.

We then investigated the role of physical activity on the profile of circulating myeloid and lymphoid immune cells in sedentary and physically active WT mice. Using cell surface markers (Supp. Table 1), we identified monocytes, neutrophils, natural killer (NK) cells, B cells, dendritic cells, as well as CD4+ and CD8+ T cells. Our results showed that the only significantly increased circulating immune cells in our mouse model were neutrophils (Figure 1F and Supplementary Figure 1). Using a cell-based assay for cytokine detection, we interrogated the serum cytokine levels of SED and PA mice and found that CCL5, which increases neutrophil accumulation, and IL-1a, which is produced by neutrophils, were both significantly increased in PA mice following 12 weeks of physical activity (Figure 1G). Conversely, CXCL10 levels showed no difference between SED and PA mice.

Given our findings on the impact of physical activity on HSCs, we asked whether the increase in circulating neutrophil levels was dictated at the HSC level. We isolated HSCs from SED and PA mice and performed an *in vitro* differentiation assay towards neutrophils. Our results show that PA HSCs have an increased rate of differentiation towards neutrophils (p=0.05) (Figure 1H, I). Together, these results show that physical activity leads to increased levels of circulating neutrophils, via preferential differentiation of HSCs towards neutrophils, as well as via cytokines that lead to neutrophil proliferation.

### Physical activity regulates the immune response in a murine model of breast cancer

Physical activity is associated with a decreased risk for breast cancer development^5, 35^, although the mechanisms for the protective effects of physical activity against breast cancer development remain largely unknown^25^. We hypothesized that physical activity exerts its protective effect on breast cancer development by reducing the oxidative stress in hematopoietic stem cells, which then differentiate into more potent tumor-infiltrating immune cells that eventually leads to a reduction in tumor incidence and/or severity (Figure 2A). In order to analyze spontaneously occurring murine breast cancer, we used the MMTV-PyMT (MP) genetically engineered mouse model; these mice are negative for both the estrogen and progesterone receptors, as well as HER2 negative^20, 29, 48^. By 3.5 months of age, they develop lung metastasis and immune cell infiltration into tumors^56^. Female mice were weaned into social housing with access to voluntary running wheels (MP PA) and aged-matched mice were weaned into regular cages with no access to running wheels, thereby constituting the sedentary group (MP SED). Daily running distance was monitored, and the MP mice were recorded to run approximately 4.4 ± 1.0 km per night. Tumor number and size were assessed for 12, 15 and 18 weeks via caliper measurement, and tumor weights were measured at the time of sacrifice. At 12 weeks of age, MP PA mice had a 40% decrease in tumor incidence (Figure 2B), but no changes in tumor weight (Figure 2C) nor tumor volume (Figure 2D). In contrast, at 15 weeks of age, MP PA mice had a 10% decrease in tumor incidence (Figure 2B), and no significant changes in tumor weight or tumor volume (Figures 2C, D). Finally, 18-week-old MP PA mice had a 10% decrease in tumor incidence (Figure 2B), a 40% decrease in tumor weight, and a significant 63% decrease in tumor volume (Figures 2C, D). Taken together, these results indicate that the impact of physical activity on tumor incidence is most significant at the early stage of tumor development, and that the impact on tumor size is most significant at the later stages of tumor development in a spontaneous model of triple negative breast cancer.

**Figure 2.**
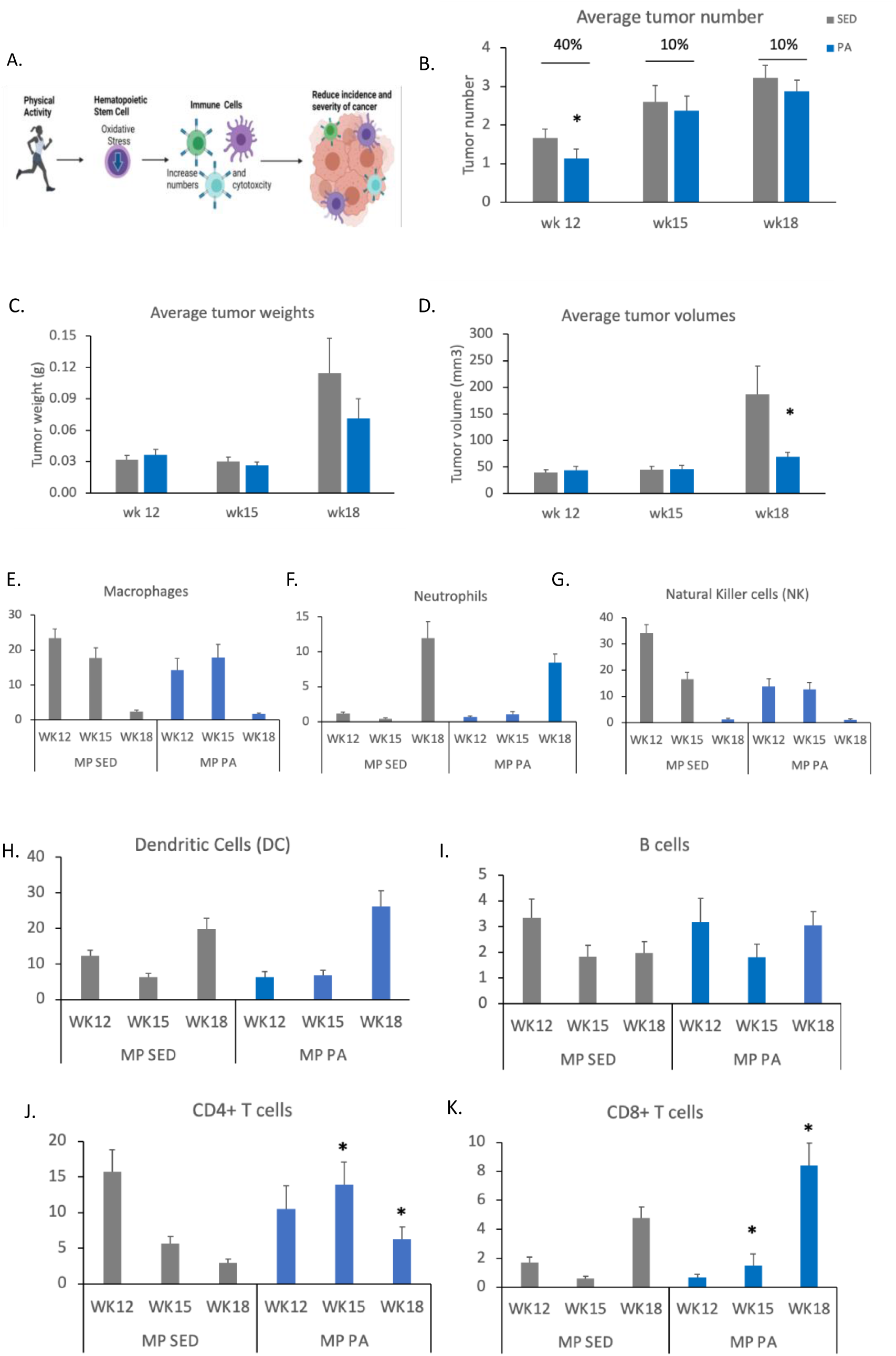
Physical activity regulates the immune response in a murine model of breast cancer. **A.** Schematic of study hypothesis (Generated with Biorender). **B.** Tumor incidence in 12-, 15- and 18-week-old SED and PA MMTV-PyMT (MP) mice. **C.** Average tumor weights in 12-, 15- and 18-week-old MP SED and PA mice. **D.** Average tumor volumes in 12-, 15- and 18-week-old MP SED and PA mice. (MP SED mice, n=23, 10 and 9; MP PA mice, n=28, 11 and 8; SED tumors, n= 42, 26 and 29; PA tumors, n= 30, 26 and 23; at 12, 15 and 18 weeks, respectively). **E-K.** Quantification of flow cytometry analysis of tumor-infiltrating immune cells from 12-, 15- and 18-week-old MP SED and MP PA mice, as a % of CD45+ cells. **E.** Macrophages (CD45+ Ly6C+F4/80+)). **F.** Neutrophils (CD45+CD11c-Ly6C+Ly6G+). **G.** Natural Killer (NK) (CD45+CD49b+NKp46+). **H.** Dendritic cells (DC) (CD45+CD11c+MHCII+). **I.** B cells (CD45+TCRb-CD19+). **J.** CD4+ T cells (CD45+TCRb+CD4+). **K.** CD8+ T cells (CD45+TCRb+CD8+). (Error bars=SEM; *, p,0.05; 2-tailed Student’s t-test).

A prevailing concept is that physical activity reduces cancer risk via direct effects on immune cells to improve cancer immunosurveillance^8, 9^. We surveyed myeloid and lymphoid circulating immune cells at 9, 12, 15 and 18 weeks of age and found no significant changes in their frequency in MP PA mice compared to MP SED (Supp. Figure 2). We then examined myeloid and lymphoid tumor-infiltrating immune cells (Figures 2E-K). We did not observe a significant effect of physical activity on tumor-infiltrating macrophages (Figure 2E), neutrophils (Figure 2F), dendritic cells (Figure 2H) or B cells (Figure 2I). Natural Killer (NK) cells were significantly decreased in the tumors of 12-week-old physically active mice (Figures 2G). Interestingly, when we examined tumor infiltrating T cells, we found that both CD4+ and CD8+ T cells were significantly increased in the tumors of physically active mice at 15 and 18 weeks of age (Figures 2J, K). Indeed, in PA mice, intra-tumoral CD4+ T cells were increased 2.5-fold and 2-fold, and intra-tumoral CD8+ T cells were increased 2.5-fold and 1.8-fold in 15-week-old and 18-week-old mice, respectively (Figures 2J, K). These results show that spontaneous physical activity in an orthotopic model of breast cancer is sufficient to induce a decrease in tumor incidence at the early stage of tumor development. Furthermore, physical activity also leads to an increase in T helper CD4+ and T cytotoxic CD8+ T cells in the tumor microenvironment, which has been shown to be a positive prognostic factor in triple negative breast cancer^31^.

### Physical activity regulates the immune response in a murine model of breast cancer via a hematopoietic stem cell-autonomous mechanism

Based on our previous findings^55, 59^, we decided to determine whether physical activity reduces the incidence and pathological severity of breast cancer by regulating immune cell number and cytotoxicity in a breast cancer tumor through reprogramming in HSCs.

HSCs were isolated from the bone marrow of PA and SED WT mice. PA mice were weaned into social housing with access to voluntary running wheels for 12 weeks; age-matched sedentary donors were used as controls. Breast cancer was studied in three groups: sedentary MP mice, sedentary MP mice reconstituted with HSCs from sedentary mice (SED→ SED), and sedentary MP mice reconstituted with HSCs from PA mice (PA→ SED) (Figure 3A). Transplantation of PA HSCs (3×10^3^ cells in 100 μl PBS into the retro-orbital plexus) into lethally irradiated MP SED recipients allows for determination of the long-term effects of PA HSCs to reduce the incidence and pathological severity of breast cancer. As we reported for HSCs derived from WT PA mice (Figure 1B), MP PA HSCs had a significant decrease in oxidant stress, as measured by a DCF-DA assay (Figure 3B). This model allowed for a thorough investigation of the changes in the tumor microenvironment as well as for the possibility to study tumor occurrence at early time points, as breast cancer tumors develop spontaneously in the MP mice. The MP SED and SED→ SED mice were used to control for the effects of irradiation and transplantation. The reconstitution capacity of the HSCs from SED and PA mice into MP SED recipient mice was similar (2-3% of total bone marrow population). This was assessed by staining of HSCs and analysis by flow cytometry prior to transplantation, as well as following 6-8 weeks of engraftment into lethally irradiated MP SED recipients. Tumor number and size were assessed in 12-15-week-old mice by caliper measurement and tumor weights were measured at the time of sacrifice. MP SED mice transplanted with HSCs from PA mice had a 26% decrease in tumor incidence (Figure 3C), closely recapitulating the decrease in tumor incidence in MP PA mice. No changes in tumor weight nor tumor volume were observed in PA→ SED mice (Supplementary Figure 2A, B).

**Figure 3.**
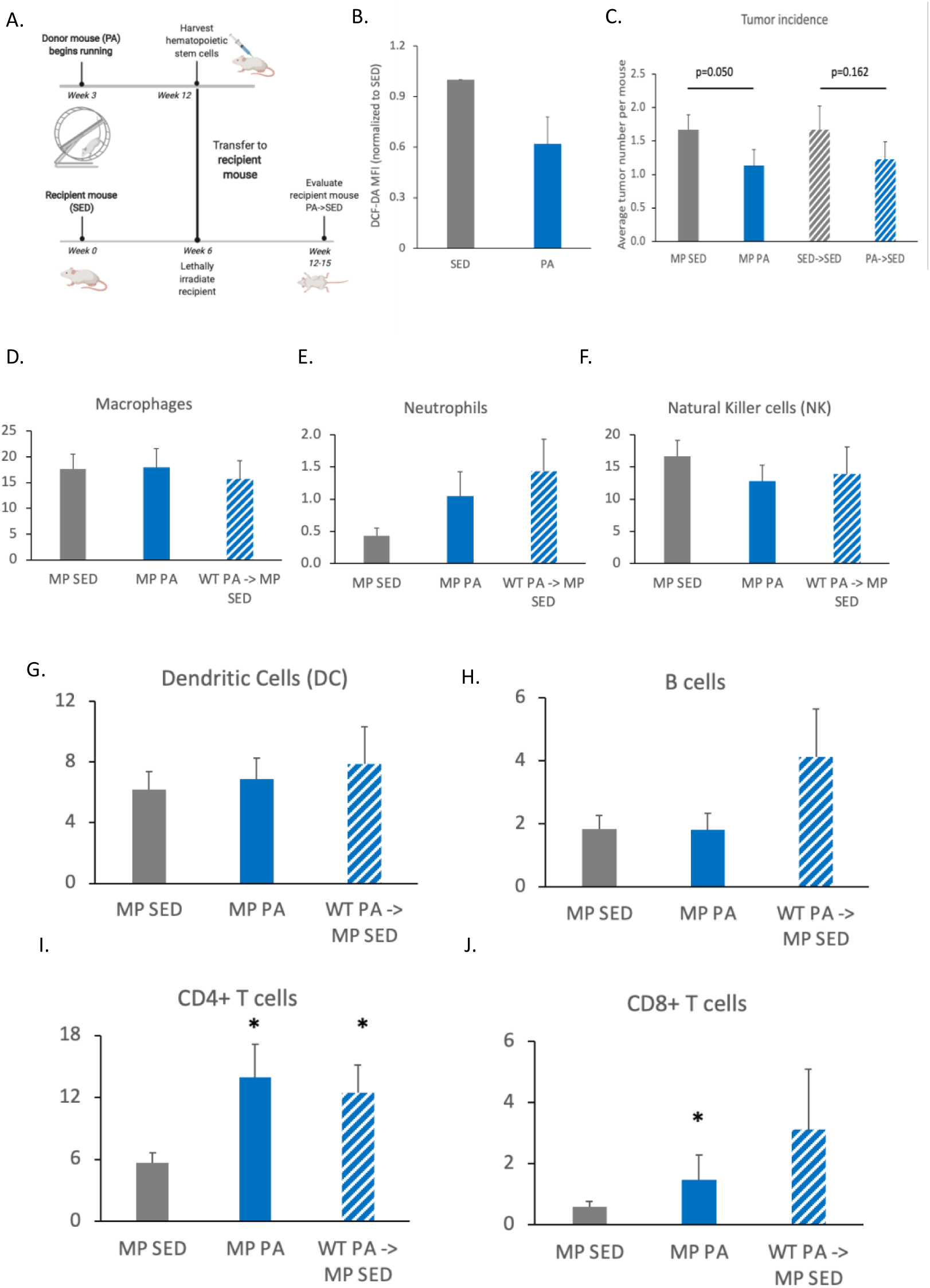
Physical activity regulates the immune response in a murine model of breast cancer via a hematopoietic stem cell-autonomous mechanism. **A.** Schematic of experimental design (Generated with Biorender). **B.** Quantification of the DCF-DA mean fluorescence intensity (MFI) in HSCs from MP SED and PA mice (n=5). **C.** Tumor incidence in 12-week-old MP SED, MP PA, SED→ SED and PA→ SED mice. (MP SED mice, n=23; MP PA mice, n=28; SED→ SED mice, n=12; PA→ SED mice, n=13; SED tumors, n= 42; PA tumors, n= 30; SED→ SED tumors, n=20; PA→ SED tumors n=16). **D-J.** Quantification of flow cytometry analysis of tumor-infiltrating immune cells from, MP SED, MP PA and PA→ SED mice, as a % of CD45+ cells. **D.** Macrophages (CD45+ Ly6C+F4/80+). **E.** Neutrophils (CD45+CD11c-Ly6C+Ly6G+). **F.** Natural Killer (NK) (CD45+CD49b+NKp46+). **G.** Dendritic cells (DC) (CD45+CD11c+MHCII+). **H.** B cells (CD45+TCRb-CD19+). **I.** CD4+ T cells (CD45+TCRb+CD4+). **J.** CD8+ T cells (CD45+TCRb+CD8+). (Error bars=SEM; *, p,0.05; 2-tailed Student’s t-test).

Next, we surveyed myeloid and lymphoid circulating immune cells and found no significant changes in the frequency of monocytes, neutrophils, NK cells, B cells, CD4+ and CD8+ T cells in PA→ SED mice compared to MP PA mice (Supplementary Figure 2C, D, E, G, H, I). However, we recorded a significant increase in circulating dendritic cells in PA→ SED mice (Supplementary Figure 2F). We then examined myeloid and lymphoid tumor-infiltrating immune cells in the tumors of PA→ SED mice, compared to MP PA and MP SED mice (Figures 3D-J). We did not observe a significant effect of physical activity on tumor-infiltrating macrophages (Figure 3D), neutrophils (Figure 3E), NK cells (Figure 3F), dendritic cells (Figure 3G), B cells (Figure 3H) or CD8+ T cells (Figure 3J). Interestingly, when we examined tumor infiltrating CD4+ T cells in PA→ SED mice, we found that they were significantly increased when compared to MP SED mice; and strikingly, identical to the rate observed in MP PA mice (Figures 3I). Taken together, these results show that, by transplanting sedentary mice with HSCs from a physically active mouse (PA→ SED), we are able to recapitulate the decrease in tumor incidence as well as the tumor-infiltrating phenotype observed in a physically active mouse (MP PA). This indicates that physical activity can regulate the immune response in a murine model of breast cancer via a hematopoietic stem cell-autonomous mechanism.

### Physical activity modulates gene expression in murine breast cancer tumors

Based on the differences in the effects of physical activity on tumor incidence and tumor-infiltrating immune cells at the analyzed time points, we decided to interrogate the gene expression profile of tumors collected from MP SED and MP PA mice at the early (week 12) and late (week 18) stages of tumor development and performed bulk RNASeq.

By using principal component analysis (PCA) we observed hierarchical clustering by physical activity status when evaluating the differential gene expression between SED and PA tumors collected from 12-week-old mice (Figure 4A). Similar segregated clustering of cells derived from SED and PA tumors from 18-week-old mice through PCA was also observed (Figure 4E). Tumor samples from SED mice clustered more closely together than with tumors from PA mice, which were more variable, in both 12- and 18-week-old mice (Figures 4A and E), suggesting that physical activity may be able to affect the gene expression profile in tumors in multiple ways.

**Figure 4.**
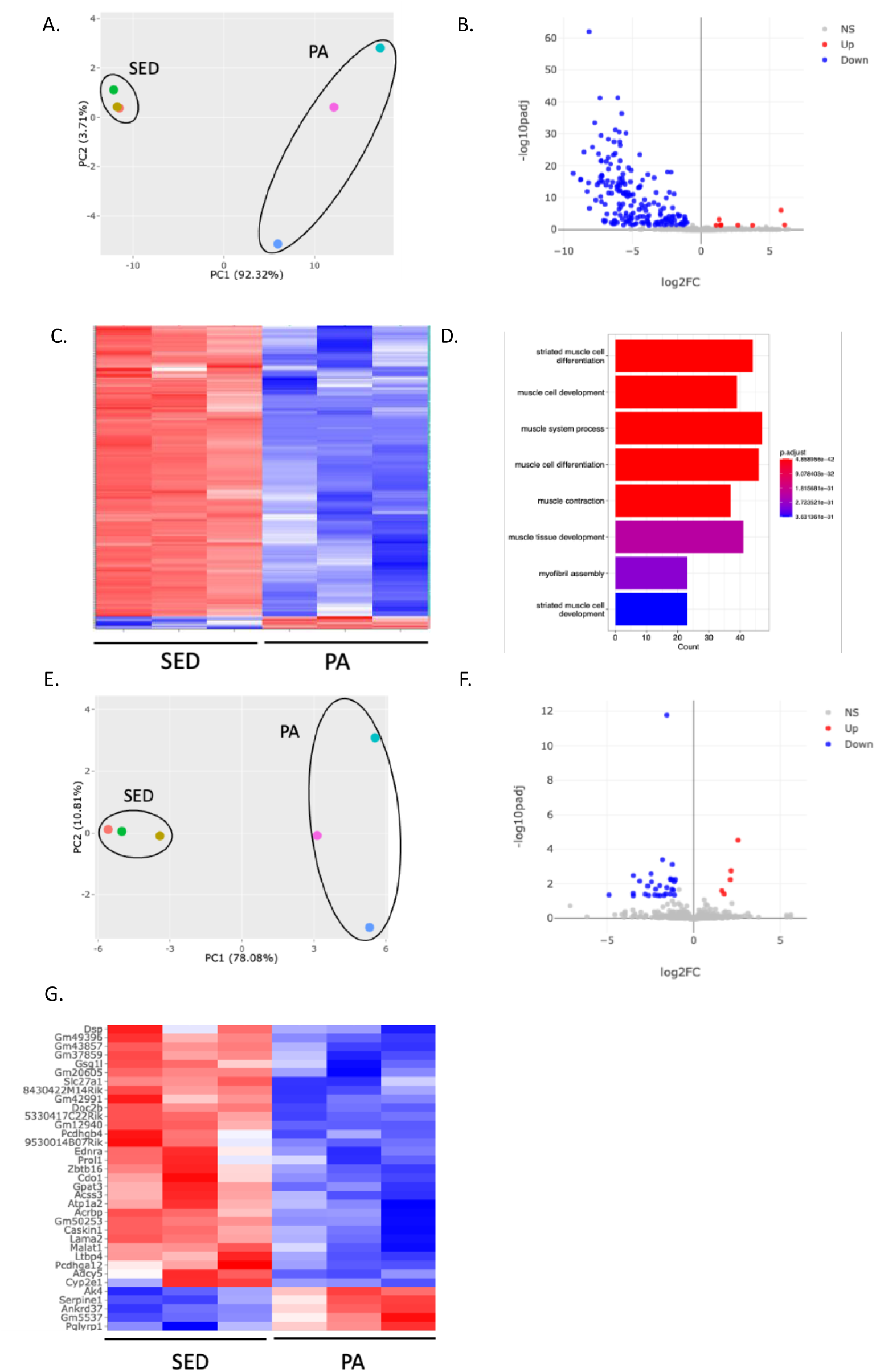
Physical activity modulates gene expression in murine breast cancer tumors. **A.** Principal component analysis (PCA) was performed on differentially expressed genes in tumors from SED and PA 12-week-old mice (n=3 tumors per condition). **B.** Volcano plot of differentially expressed genes in tumors from SED and PA 12-week-old mice. **C.** Heatmap representative of 186 downregulated and 8 upregulated genes in tumors of 12-week-old PA mice. **D**. Pathway enrichment analysis of differentially expressed genes in PA 12-week-old mice tumors, p value cut off <0.05 and fold change >2. **E.** Principal component analysis (PCA) was performed on differentially expressed genes in tumors from SED and PA 18-week-old mice (n=3 tumors per condition). **F.** Volcano plot of differentially expressed genes in murine tumors from SED and PA 18-week-old mice. **G.** Heatmap representative of 30 downregulated and 5 upregulated genes in tumors of 18-week-old PA mice, p value cut off <0.05 and fold change >2.

We identified 8 upregulated genes and 186 downregulated genes in tumors from 12-week-old PA mice (padj=0.05, log2FC =2) (Figure 4B, C). Pathway enrichment analysis for differentially expressed genes showed a significant enrichment in muscle cell differentiation, muscle system process, muscle cell development, muscle tissue development, and myofibril assembly pathways (Figure 4D) in tumors of 12-week-old SED mice. While intriguing, these results point to a significant negative effect of physical activity on the levels of myoepithelial genes in tumors, which have been identified as poor prognosticators for breast cancer development and metastasis^11, 13, 39^. Furthermore, the upregulated genes in MP PA tumors included members of the Homeobox family, *Hoxb2*, *Hoxb3* and *Hoxc10*. *Hoxb2* has been reported to act as a tumor suppressor gene in murine breast cancer^4^, and loss of *Hoxb3* correlates with the development of hormone receptor negative breast cancer^62^. Therefore, increased levels of these two genes in the tumors of physically active mice may explain the reduced incidence phenotype we have observed in 12-week-old PA mice.

When we performed a similar analysis on tumors isolated from 18-week-old mice, we identified 5 upregulated genes and 30 downregulated genes in MP PA tumors (padj=0.05, log2FC=2) (Figure 4F, G) and no significant pathway enrichment was detected, most likely due to the reduced number of differentially expressed genes. Of note, the upregulation of *Pglyrp1*, which has been shown to activate CD4+ T cells^44^, and the downregulation of *Slc27a*, *Lama2* and *Malat1*, which have been identified as poor prognosis markers in several breast cancer models^16,27, 33, 60^.

### Physical activity modulates gene expression in HSCs

Given the effect of physical activity on HSC oxidant stress, and our identification of a novel role for HSCs in the regulation of the immune response in triple negative breast cancer, we decided to investigate gene expression profiles in HSCs from MP SED and PA mice, using bulk RNASeq. By using principal component analysis (PCA) we observed hierarchical clustering by physical activity status when evaluating the differential gene expression between SED and PA HSCs collected from 12-week-old mice (Figure 5A). HSCs from SED mice clustered more closely together than with HSCs from PA mice, which were more variable, (Figures 5A), suggesting that physical activity may be able to affect the gene expression profile in HSCs in multiple ways, much like what we observed in our tumor analyses (Figure 4A, E). We identified 186 upregulated genes and 2 downregulated genes in HSCs from 12-week-old PA mice (Figure 5B, C) (For Figure 5B: padj=0.05, log2FC=2; for Figure 5C: padj=0.01, log2FC=2). Pathway enrichment analysis for differentially expressed genes showed a significant enrichment in myeloid leukocyte activation, leukocyte migration and chemotaxis, blood coagulation and hemostasis pathways (Figure 5D) in HSCs of 12-week-old PA mice. One of the two downregulated genes in PA HSCs, *Uchl1*, has been recently identified as a novel target in triple negative breast cancer^34^. Understanding the mechanisms driven by *Uchl1* in HSCs may uncover novel therapeutic approaches.

**Figure 5.**
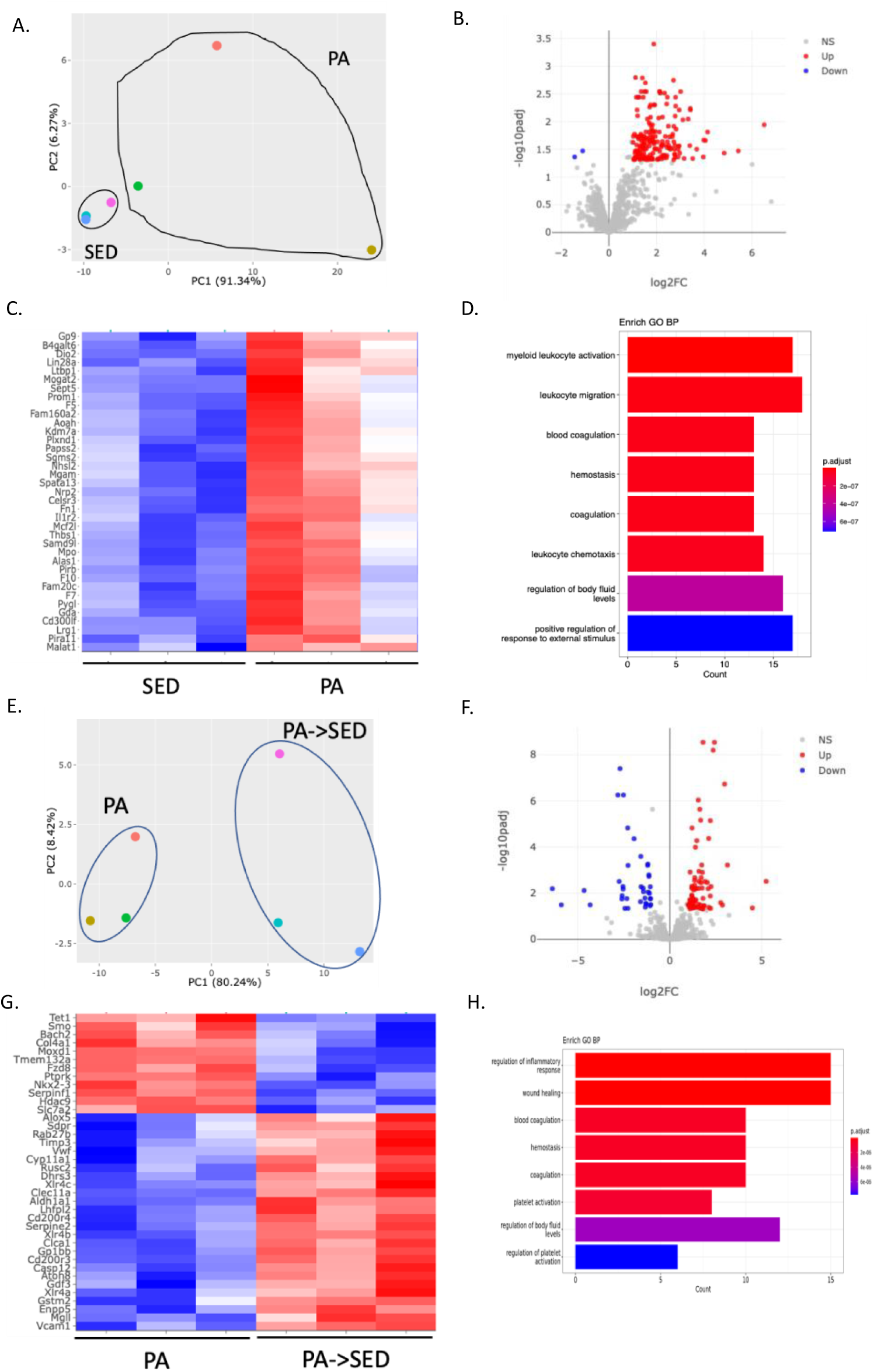
Physical activity modulates gene expression in HSCs. **A.** Principal component analysis (PCA) was performed on differentially expressed genes in HSCs from SED and PA mice (n=3 samples per condition). **B.** Volcano plot of differentially expressed genes in HSCs from PA and PA→ SED mice. **C.** Heatmap representative of 37 upregulated genes in HSCs of PA mice, p value cut off <0.01 and fold change >2. **D**. Pathway enrichment analysis of differentially expressed genes in HSCs from SED and PA mice, p value cut off <0.05 and fold change >2. **E.** Principal component analysis (PCA) was performed on differentially expressed genes in HSCs from PA and PA→ SED mice (n=3 samples per condition). **F.** Volcano plot of differentially expressed genes in HSCs from PA and PA→ SED mice. **G.** Heatmap representative of 26 upregulated genes and 12 downregulated genes in HSCs from PA→ SED mice, p value cut off <0.005 and fold change >2. **H**. Pathway enrichment analysis of differentially expressed genes in PA→ SED mice, p value cut off <0.05 and fold change >2.

Finally, given the similarities in the tumor incidence phenotype as well as in the CD4+ T cells intra-tumoral infiltration in MP PA and PA→ SED mice, we compared the gene expression profile of PA and PA→ SED HSCs by bulk RNASeq. Principal component analysis (PCA) showed hierarchical clustering when evaluating the differential gene expression between PA and PA→ SED HSCs collected from 12-15-week-old mice (Figure 5E). We identified 78 upregulated genes and 49 downregulated genes in PA→ SED HSCs (Figure 5F, G) (For Figure 5F: padj=0.05, log2FC=2; for Figure 5G: padj=0.005, log2FC=2). Pathway enrichment analysis for differentially expressed genes showed a significant enrichment in the regulation of the inflammatory response, blood coagulation, platelet activation and hemostasis pathways (Figure 5H) in HSCs of PA→ SED mice, similarly to what we observed for HSCs from MP PA mice (Figure 5D). Among the downregulated genes in PA→ SED HSCs are *Smo, Col4a1, Ptprk* and *Tet1*, which have been associated with advanced tumor stage, proliferation and migration of ductal carcinomas, poor prognosis and activation of oncogenic signaling in triple negative breast cancer^15, 22, 53, 54^.

## DISCUSSION

In this study, we show that physical activity exerts its protective effect on breast cancer development by reducing the oxidative stress and regulating the gene expression in hematopoietic stem cells. This leads to an imprinting of the HSCs, whereby sedentary mice transplanted with HSCs from physically active mice display an identical increase in tumor infiltrating CD4+ T cells to that observed in physically active mice. This increase in the rate of tumor-infiltrating immune cells leads to a reduction in tumor incidence at early stages of tumor development and a reduction in tumor volume at late stages of tumor development. Furthermore, in a pilot study in NSG mice transplanted with tumor cells from a triple negative breast cancer patient, we observed a similar decrease in tumor volume, following 4 weeks of physical activity, compared to transplanted NSG mice that were sedentary (Supp. Figure 3A).

In the analyses of differentially expressed genes in HSCs from MP PA and PA→ SED mice, we identified similar enrichment in immune and inflammatory pathways, indicating that the changes in gene expression in HSCs induced by physical activity are maintained even when the cells are engrafted in a sedentary animal. Furthermore, we identified several epigenetic regulators that were significantly dysregulated in HSCs. Indeed, *Kdm7a* is significantly upregulated in MP PA HSCs and *Tet1* and *Hdac9* are significantly downregulated in PA→ SED HSCs. These results point to a potential role for these regulators in the imprinting of HSCs, which may dictate their differentiation towards cytotoxic tumor infiltrating immune cells.

Acute exercise may affect HSC number by influencing HSC mobilization and bone marrow HSC proliferation. However, HSC function is potentially more important than their number. To date, very few studies have investigated the effect of an acute bout of exercise on HSC function. Kröpfl et al.^52^ showed that an ultra-endurance cycling race mobilized HSCs, but that HSC function, measured via colony forming capacity *in vitro*, was decreased after the strenuous activity in humans. Furthermore, they determined that the decrease in HSC function was correlated to an increase in oxidative stress. Another study also showed that a single bout of relatively intense cycle ergometry increased the numbers of circulating hematopoietic progenitors while negatively affecting their *in vitro* colony forming functionality^26^; however, mechanisms for the decrease in HSC function were not investigated. Acute bouts of exercise, especially highly strenuous or unaccustomed exercise, are known to induce oxidative stress. The extreme or unaccustomed exercise in these studies is not representative of the chronic, moderate activity that has been correlated with a decreased risk of cancer. The limited data on the effect of exercise on HSC function demonstrates that a single bout of intense or strenuous exercise has negative effects on HSC function, likely due to oxidative stress. However, the theory of exercise hormesis postulates that single bouts of exercise prime cells to defend against the oxidative damage from future oxidative insults^7^. Therefore, while a single bout of exercise may be damaging, chronic participation in moderate exercise may protect against oxidative damage in HSCs. In our model, moderate physical activity leads to a decrease in oxidative stress in both WT and MMTV-PyMT mice.

Several studies have reported the beneficial impact of physical activity on cancer incidence and severity. In a spontaneous triple negative breast cancer model, access to voluntary running wheels decreased tumor size by 40%, but was not effective at reducing the incidence of cancer^51^, potentially suggesting that the beneficial effects of physical activity on the immune system are not robust enough to overcome a strong genetic predisposition to breast cancer. In a syngeneic orthotopic transplantation model, voluntary wheel running also reduced mammary tumor volume^2^. In a chemically induced carcinoma model of breast cancer, forced treadmill running decreased tumor incidence in rats^30^. These studies highlight the heterogeneity in the response to exercise for various models of breast cancer; however, they largely fail to establish a mechanism for the cancer protective effects of regular physical activity. In our model, mice have access to running wheels at weaning, thereby mimicking the benefits of regular physical activity.

An important benefit of physical activity is its role in the stimulation of T cell expansion. The characteristics of exercise-induced leukocytosis may vary depending on body composition, but not on training status. Ndon and colleagues^36^ provided evidence that chronic training does not affect leukocytosis in response to an acute bout of exercise. In support of this, habitually active participants with high fitness and sedentary participants with low fitness had relatively similar exercise-induced leukocytosis responses, except sedentary participants had more CD4^+^CD45RO^+-^ memory T cells and CD4^+^PD-1^+^ T cells than self-reported active individuals. Leukocyte populations in the blood may be affected by other indicators of regular exercise, including lean body mass and body fat percentage. Resting levels of CD8^+^ T cells and Tregs were found to be positively related to body mass, indicating greater CD8^+^ T cells and Tregs in individuals with higher body fat percentages. Effector memory T cells were inversely related to body mass index at rest^18^. Overall, these studies demonstrate that leukocytosis occurs following physical activity in both sedentary and active individuals; however, chronic physical activity may affect the leukocytosis profile. It is unknown how the differential leukocytosis in varying populations affect the immunosurveillance ability of leukocytes from active and inactive people. In our study, we show an increase in tumor infiltrating CD4+ and CD8+ T cells following 15-18 weeks of physical activity, in a triple negative model of murine breast cancer.

Finally, using differential gene expression analyses in early and late stage tumors from MP SED and MP PA mice, we have identified an increase in tumor-suppressor genes, such as *Hoxb2* and *Pglyrp1*^4, 44^, and a decrease in potential oncogenes, such as *Slc27a1* and *Malat1*^16, 33, 60^ in the tumors of MP PA mice. Interestingly, in our pilot study in NSG mice transplanted with tumor cells from a triple negative breast cancer patient, we observed a similar decrease in the expression of *MALAT1* in the tumors isolated from NSG-PDX mice following 4 weeks of physical activity (Supp. Figure 3B). Taken together, our results may lead to the identification of potential diagnostic biomarkers, which could be utilized in future clinical and translational studies to engineer immunotherapies for breast cancer patients. Our results may be further exploited in the usage of physical activity as an adjuvant therapy to tune the immune system of patients undergoing conventional treatments.

## MATERIALS AND METHODS

### Animals

Female C57BL/6J (strain number: 000664), male MMTV-PyMT (strain number: 022974) and female NOD.*Cg-Prkdc^scid^Il2rg^tm1Wjl^/SzJ* (NOD-*scid IL2rγ^null^*, NSG) strain number: 005557) mice were purchased from The Jackson Laboratory (Bar Harbor, ME), fed a standard chow (5.4 g fat/100 g diet), and maintained on a 12-hour light/dark cycle. Pups resulting from breeding female C57BL/6J with male MMTV-PyMT were genotyped following the strain-specific protocol, listed on The Jackson Laboratory website; female mice carrying the MMTV-PyMT gene were selected for experiments. Care was in accordance with NIH guidelines. All mice were housed socially with 2-4 mice per cage. All protocols were approved by the Institutional Animal Care and Use Committee at the University of Massachusetts Medical School.

### PDX tumor explants

Patient tumor explants were obtained from surgical specimens of breast cancer from patients at the UMass Memorial Hospital (Worcester, MA, USA). Written, informed consent was obtained from the patients before collection of specimens. PDX models were generated by implantation of PDX into NSG mice. In brief, patient-derived tumors were finely minced and loaded into 1-cc syringes with 14-gauge needles. 20–40 μl of homogenized tumor tissue was inoculated subcutaneously at the right flank of NSG mice while under anesthesia. Tumors were measured by caliper every 7 days, and volumes (mm^3^) were calculated by (length × width)^2^/2^58^.

### Wheel running

At four weeks of age, MMTV-PyMT female mice in the physically active group were given a voluntary running wheel (Columbus Instruments, Columbus, OH), and age-matched sedentary mice were maintained in cages without running wheels. NSG-PDX mice were transferred to running wheel-fitted cages the day following tumor cell injection. A magnetic counter was mounted to the outside of the cages, with a wire connected to a computer to record wheel revolutions 24 hours a day, 7 days a week. The amount of voluntary wheel running was continuously monitored and quantified by multiplying the number of wheel revolutions by the circumference of the running wheel. An estimation of the distance run per mouse was calculated by dividing the total distance by the number of the mice in the cage, as previously reported^6^.

### Hematopoietic stem cell isolation

HSCs were isolated using fluorescence activated cell sorting (FACS). Briefly, mice were sacrificed with an overdose of isoflurane followed by cervical dislocation. Whole bone marrow was flushed from the tibias and femurs, centrifuged, and resuspended in FACS buffer containing 1% FBS and 1% penicillin/streptomycin. Next, HSCs were enriched using the EasySep™ Mouse Hematopoietic Progenitor Cell Isolation Kit (STEMCELL Technologies Inc) according to manufacturer’s instructions. Briefly, cells were incubated with rat serum to block non-specific binding and a biotinylated antibody against lineage positive, non-hematopoietic stem or progenitor cells. Streptavidin-coated magnetic particles were incubated with the cells, and lineage positive labeled cells were separated using a magnet. The enriched cell population was poured off and stained with fluorescently-conjugated monoclonal antibodies against cKit, Sca1, CD90.1, and a lineage cocktail consisting of CD4, CD8, B220, TER-119, Mac-1, and Gr-1. FACS was used to sort HSCs, which were defined as cKit^+^Sca-1^+^CD90.1^lo/-^Lin^-^.

### Leukocyte and intra-tumoral immunophenotyping

Cheek punch was used to collect peripheral blood from mice. Blood was collected into tubes containing heparin, and RBC lysis was performed using an ammonium chloride RBC lysis buffer (Boston Bioproducts; cat# IBB-197X). Tumors were excised from mouse mammary pads at specified time points and minced to 1mm fragments in 1ml cold PBS on ice. They were subsequently incubated with Liberase-TL (Sigma; cat# 5401020001), Liberase-DL (Sigma; cat# 5401160001), and DNAseI (Sigma; cat#10104159001) for 1 hour at 37C on a rotating platform. Following enzymatic digestion, samples were filtered through a 70um filter, washed with FACS buffer and RBC lysis was performed using an ammonium chloride RBC lysis buffer (Boston Bioproducts; cat# IBB-197X).

Following lysis, cells were stained with monoclonal antibodies directed against specific cell surface epitopes as detailed in Tables. All antibodies had previously been titrated and were used at a concentration where the mean fluorescent intensity plateaus. The antibodies used and the identification of immune cell populations are given in Tables. Cells were incubated with the antibody mix in FACS buffer containing 1% FBS and 1% penicillin/streptomycin for 20 min. at 4 C, washed with FACS buffer and then re-suspended in PBS containing Zombie Green (Biolegend; cat#423111) following manufacturer’s recommendation.

After surface staining, cells were analyzed by flow cytometry using either an Aurora Cytek (Cytek Bio) equipped with 350 nm, 405 nm, 488 nm, 561 nm and 641 nm excitation lasers. Prior to the analysis of cells, compensation was manually adjusted using UltraComp eBeads (eBioscience; cat# 01-2222-42) stained with single antibodies. Analysis of flow cytometric data was performed using FlowJo software (Tree Star). Populations were gated according to the markers listed in supplemental information.

### Irradiation of Transplant Recipients and HSC transplantation

Recipient mice were irradiated (two doses of 550 rad, 6 hours apart, using 137Cs irradiator in the ASC mouse barrier facility). Mice were then placed under general anesthesia utilizing 3% isoflurane delivered in 100% O2 using a nose cone and placed on a warm pad over a circulating water blanket. One drop of Proparacaine HCL (0.5%) was applied to each eye for local anesthesia, followed by a sterile saline rinse. 3000 HSCs in 100 μl sterile PBS were injected via the retro-orbital plexus as a single injection using a sterile disposable 1 ml syringe and needle. A solution of sulfamethoxazole and trimethoprim (Septra) (157 mg/kg sulfamethoxazole and 33mg/kg trimethoprim) was added to the drinking water for 2 weeks post-transplant for antibiotic prophylaxis.

### Oxidative stress quantification

Freshly sorted HSCs were cultured overnight in medium supplemented to maintain stemness of HSCs (RPMI 1640 containing 20% FBS, 1% P/S, 50 ug/ml beta mercaptoethanol, 1% Glutamax, 1% nonessential amino acids, and 1% sodium pyruvate supplemented with 10 ug/ml SCF, 2 ug/ml TPO, 5 ug/ml Flt3L, 1.2 ug/ml IL-3, and 2 ug/ml IL-6 (Peprotech). To quantify oxidative stress, HSCs were harvested after overnight culture and stained with DCF (1:1000) for 30 min at 37°C. Flow cytometry was performed on a BDBiosciences LSR to measure DCF-DA fluorescence in the FITC channel of the stained samples and unstained negative control samples. Quantification was performed in FlowJo as median fluorescence intensity (MFI) of the positively stained cells relative to the unstained cells.

### Gene expression

RNA was isolated from freshly sorted HSCs using the RNAqueous™-Micro Total RNA Isolation Kit (Invitrogen) according to manufacturer’s instructions. cDNA was synthesize using the SuperScriptTM III First-Strand Synthesis SuperMix for qRT-PCR (Invitrogen). The expression of genes related to oxidative stress and antioxidant defense or epigenetic modifying enzymes in physically active compared to sedentary mice were profiled using the RT² Profiler™ PCR Array Mouse Oxidative Stress and Antioxidant Defense kit or the RT² Profiler™ PCR Array Mouse Epigenetic Chromatin Modification Enzymes kit, respectively (Qiagen). Data from the arrays were analyzed using the Qiagen Data Analysis Center to identify genes that were greater than 2.0-fold up- or downregulated in the physically active than in the sedentary mice. qRT-PCR was used to confirm dysregulated gene expression of the genes identified in the qPCR array (see table for primers). qRT-PCR was performed using Kapa SYBR FAST qPCR kit (Kapa Biosystems).

### Bulk RNA-Seq

RNA was extracted from mouse mammary tumors using the TRIzol method. Library preparation was performed using TruSeq Stranded mRNA Sample Prep Kit (Cat# 20020594, Illumina) according to manufacturer’s instruction. The libraries were sequenced on the NextSeq 500 system (Illumina) (2×150bp paired-end reads). The generated fastq files were loaded into the DolphinNext platform (https://dolphinnext.umassmed.edu/) and the Bulk RNA sequencing pipeline was used^61^. The fastq files were aligned to the mouse (mm10) genome. Once aligned, the files were run through RSEM for normalization. Differential expression analysis was performed using the DEBrowser platform (https://debrowser.umassmed.edu/). Pathway analysis was performed by using the Gene ontology analysis on the DEBrowser platform. The data discussed in this publication have been deposited in NCBI’s Gene Expression Omnibus^12^ and are accessible through GEO Series accession number GSE229575.

### HSC differentiation to neutrophils

Hematopoietic stem cells were isolated from mice, as described earlier in this section. As described in Gupta et al^17^, cells were cultured for 3 days in IMDM, 20% Horse Serum, and 50 ng/ml each of SCF and IL-3. After 3 days of culture, cells are expanded into medium with 50 ng/ml each of SCF, IL-3 and G-CSF, and incubated for an additional 2 days. The cells are then collected by centrifugation at 250 × g for 5 min, washed twice in 1× PBS, and cultured in IMDM, 20% Horse Serum and 50 ng/ml G-CSF for an additional two days, bringing the total number of days to 7 days from day of harvest.

### Cell-based assay for cytokine detection

For the detection of the cytokines, cytometric bead array (CBA) assay was performed using BioLegend LEGENDplex Mouse Inflammation Panel (BioLegend, catalog #740446) following the manufacturer’s recommendations. Serum samples from experimental animals were diluted 1:4. Data were acquired on a BD Celesta and analyzed using the LEGENDplex Data Analysis Software Suite.

## ACKNOWLEDGEMENTS

This study was supported in part by NIH grant R21AG067376 to LM and by the Joseph P. Healey Endowment to LK. We acknowledge the use of services from the following UMASS cores: Flow cytometry and Bioinformatics.

## AUTHOR CONTRIBUTIONS

LK: hypothesis generation, conceptual design, experiment design and performance, data analysis, manuscript preparation. KH: experiment design and performance, data analysis, manuscript preparation. AT: experiment performance, data analysis, manuscript preparation. AS: experiment design and performance, manuscript preparation. LF: experiment design and performance, manuscript preparation. TTN: conceptual design, manuscript preparation. MB: conceptual design, manuscript preparation. LMM: conceptual design, manuscript preparation.

