## Supplementary figures and tables for "Physical activity regulates the immune response to breast cancer by a hematopoietic stem cell-autonomous mechanism"

**Supplementary Figure 1**

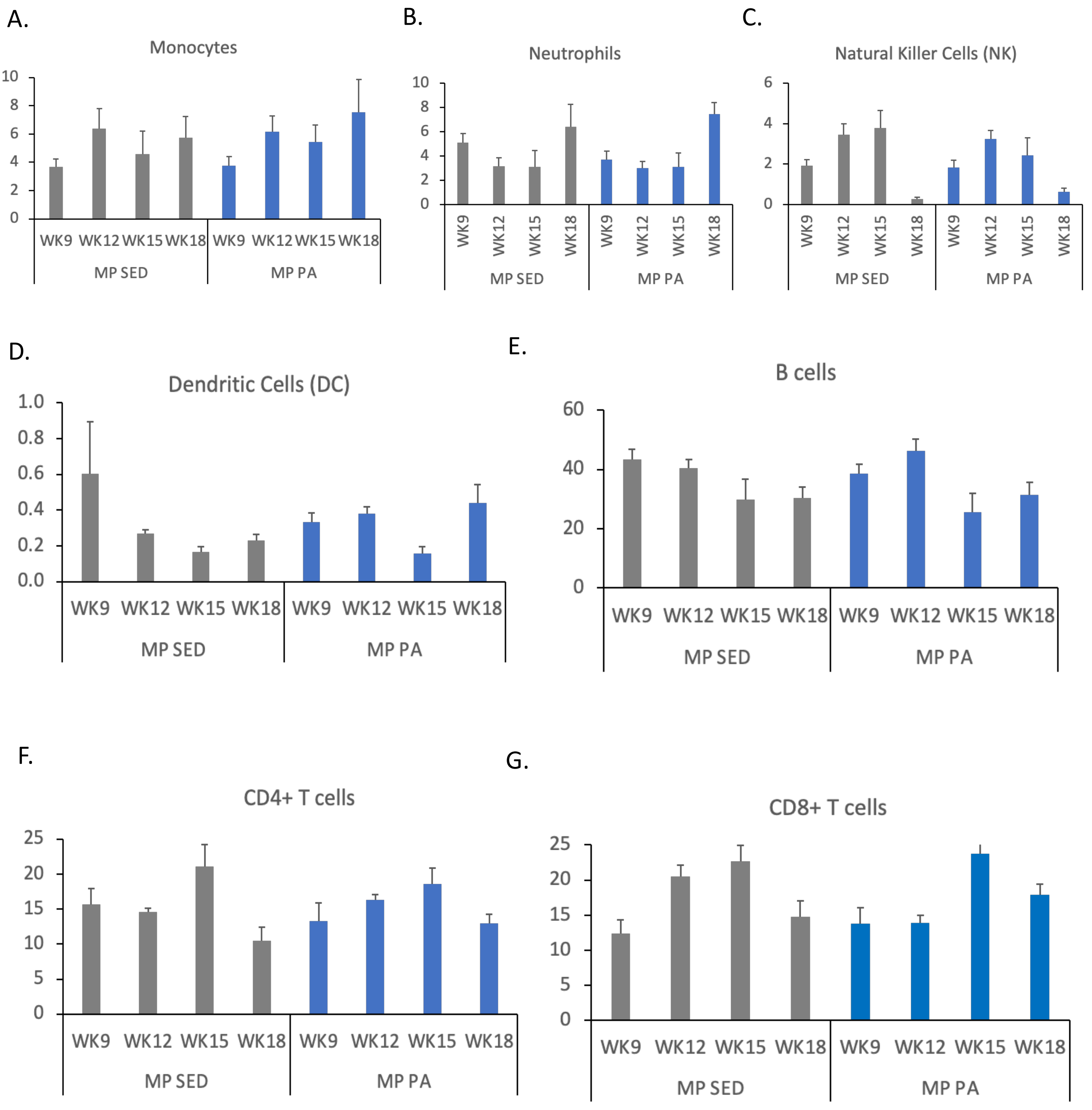


**Supplementary Figure 1. Circulating myeloid and lymphoid immune cells in 12-, 15- and 18-week-old MP SED and MP PA mice.** Quantification of flow cytometry analysis of circulating immune cells from 9-, 12-, 15- and 18-week-old MP SED and MP PA mice, as a % of CD45+ cells. **A.** Monocytes (CD45+ Ly6C+CD11b+). **B.** Neutrophils (CD45+CD11c-Ly6C+Ly6G+). **C.** Natural Killer (NK) cells (CD45+CD49b+NKp46+). **D.** Dendritic cells (DC) (CD45+CD11c+MHCII+). **E.** B cells (CD45+TCRb-CD19+). **F.** CD4+ T cells (CD45+TCRb+CD4+). **G.** CD8+ T cells (CD45+TCRb+CD8+). (MP SED mice, n=10, 23, 10 and 9; MP PA mice, n=10, 28, 11 and 8; Error bars=SEM).

**Supplementary Figure 2**

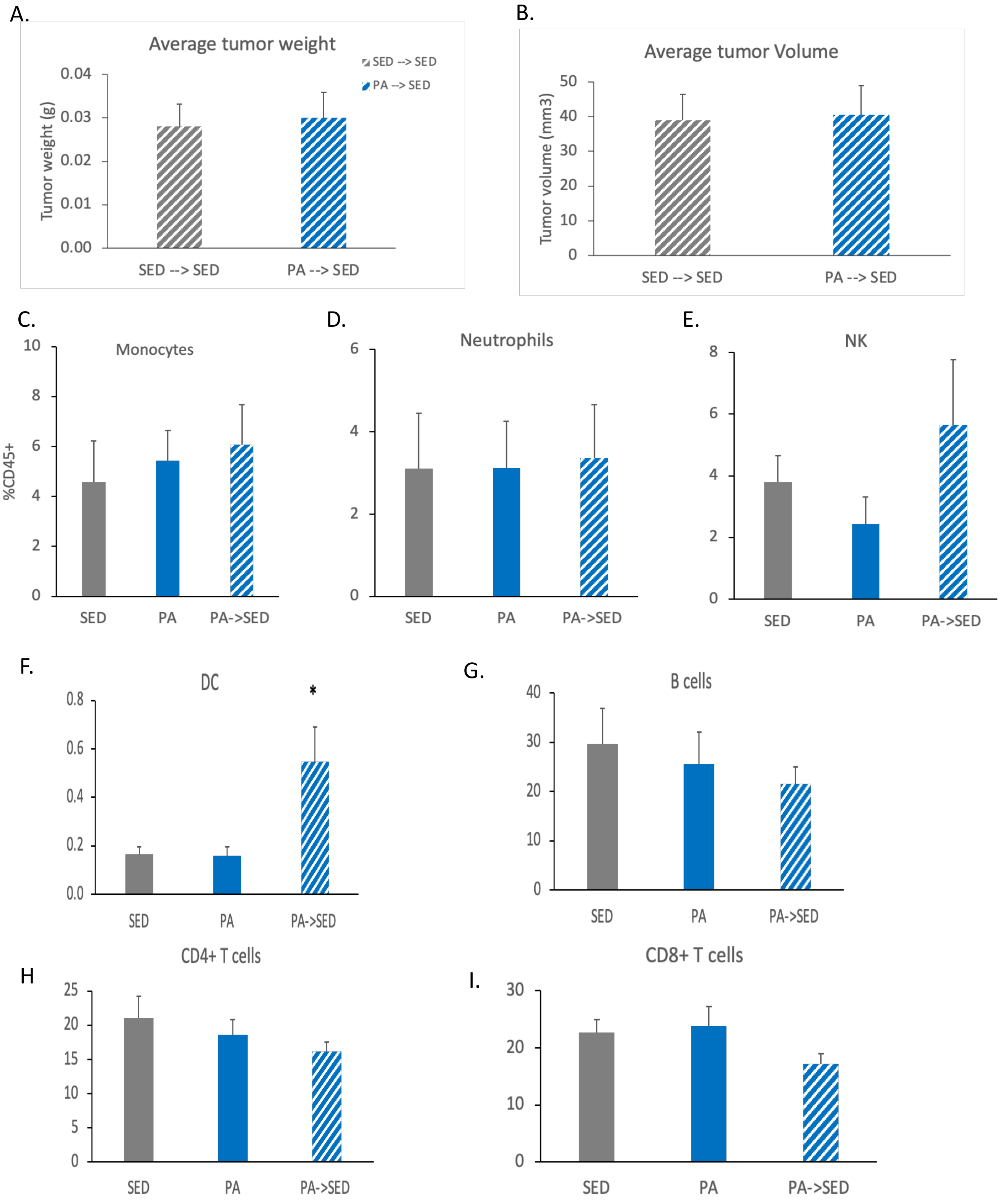


**Supplementary Figure 2. Analysis of tumors and circulating myeloid and lymphoid immune cells in PA→ SED mice. A.** Average tumor weights in MP SED, MP PA and PA→ SED mice. **B.** Average tumor volumes in MP SED, MP PA and PA→ SED mice. **C-I**. Quantification of flow cytometry analysis of circulating immune cells from MP SED, MP PA and PA→ SED mice, as a % of CD45+ cells. **C.** Monocytes (CD45+Ly6C+CD11b+). **D.** Neutrophils (CD45+CD11c-Ly6C+Ly6G+). **E.** Natural Killer (NK) cells (CD45+CD49b+NKp46+). **F.** Dendritic cells (DC) (CD45+CD11c+MHCII+). **G.** B cells (CD45+TCRb-CD19+). **H.** CD4+ T cells (CD45+TCRb+CD4+). **I.** CD8+ T cells (CD45+TCRb+CD8+). (MP SED mice, n=10; MP PA mice, n=11; PA→ SED mice, n=13; Error bars=SEM; *, $p<0.05$; 2-tailed Student's t-test).

**Supplementary Figure 3**

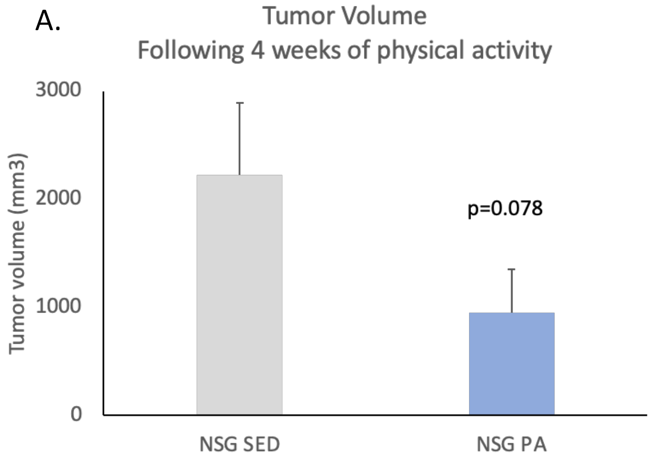


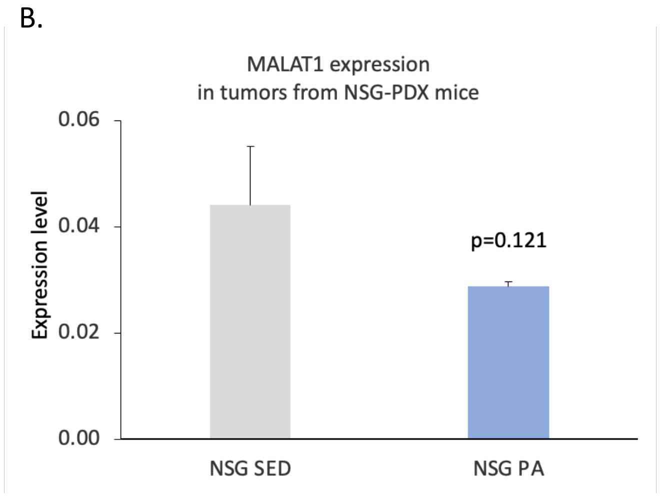


**Supplementary Figure 3. Analysis of NSG-PDX mice following 4 weeks of physical activity. A.** Average tumor volume in NSG-SED and NSG-PA mice following 4 weeks of physical activity. B. MALAT1 gene expression levels in NSG-PDX tumors from NSG-SED and NSG-PA mice. (NSG-SED mice and tumors, n=3; NSG-PA mice and tumors, n=3; Error bars=SEM; p-values calculated using 1-tailed Student's t-test).

### Supplementary Table 1

| REAGENT or RESOURCE | SOURCE | Catalog# |
|---|---|---|
| **Antibodies** | | |
| Brilliant Violet 750™ anti-mouse CD4 Antibody | Biolegend | 100467 |
| PE dazzle-594 CD49b | Biolegend | 108923 |
| PerCP/Cyanine5.5 anti-mouse CD11c | Biolegend | 117327 |
| Alexa Fluor 700™ anti-mouse CD8 Antibody | Biolegend | 100729 |
| Zombie Green™ Fixable Viability Kit | Biolegend | 423111 |
| PE/Cy7 anti-mouse Ly-6G | Biolegend | 127617 |
| Brilliant Violet 711™ anti-mouse Ly-6C | Biolegend | 128037 |
| APC anti-mouse CD335 (NKp46) | Biolegend | 137608 |
| Pacific Blue™ anti-mouse CD45.2 | Biolegend | 109820 |
| APC/Fire anti-mouse CD19 | Biolegend | 115557 |
| Brilliant Violet 650™ anti-mouse/human CD11b | Biolegend | 101259 |
| Brilliant Violet 510™ anti-mouse TCR β chain | Biolegend | 109234 |
| PE anti-mouse F4/80 | Biolegend | 123127 |
| Brilliant Violet 785™ anti-mouse MHC II Antibody | Biolegend | 107645 |
| APC Lineage antibody cocktail | Fisher Scientific | 558074 |
| Anti-Mouse CD117 PE | Fisher Scientific | 12-1171-83 |
| Anti-mouse Ly6A/E FITC | eBioscience | 11-5981-85 |
| **Chemicals and Media** | | |
| Murine Flt3L | Peprotech | 250-31L |
| Murine TPO | Peprotech | 315-14 |
| Murine SCF | Peprotech | 250-03 |
| Murine IL-3 | Peprotech | 213-13 |
| murine G-CSF | Peprotech | 250-05-10 |
| IMDM | Gibco | 12440-053 |
| PBS | Gibco | 14190250 |
| RPMI Medium 1640 | Gibco | 11875-093 |
| Horse Serum | Gibco | 26050-070 |
| Penicillin/Streptomycin | Gibco | 15140122 |
| 10mM MEM Non-Essential Amino Acids | Gibco | 11140050 |
| Glutamax | Gibco | 35050061 |
| Sodium pyruvate | Gibco | 11360070 |
| **Commercial Assays** | | |
| RNAqueous®-Micro Total RNA Isolation Kit | Invitrogen | AM1931 |
| SuperScript III First strand Synthesis Super Mix for qRT-PCR | Invitrogen | 11752-050 |
| SYBR Fast Universal master mix | Kapa Biosystems | KK4602 |
| RT2 First Strand Kit | Qiagen | 330401 |
| RT2 SYBR Green ROX qPCR Mastermix | Qiagen | 330520 |
| EasySep™ Mouse Hematopoietic Progenitor Cell Isolation Kit | Stem Cell Technologies | 19856 |
| Liberase™ TL Research Grade | Sigma | 5401020001 |
| Liberase™ DL Research Grade | Sigma | 5401160001 |
| DNAseI | Sigma | 10104159001 |
| **Deposited data** | | |
| GSE229575 | NCBI's Gene Expression Omnibus | |
| **Experimental models: Organisms/strains** | | |
| C57BL6/J | Jackson Laboratories | #000664 |
| MMTV-PyMT | Jackson Laboratories | #022974 |
| **Software and algorithms** | | |
| Flow Jo software | Tree Star | N/A |
| GraphPad prism software | GraphPad Software, Inc., SanDiego, CA | www.graphpad.com |
| Biorender | Biorender.com | N/A |
| DolphinNext | https://dolphinnext.umassmed.edu/ | N/A |

### Supplementary Table 2

| PRIMER NAME | Species | SEQUENCE |
|---|---|---|
| **Cygb** | Mouse | Forward: ACCCTCTCCACATAGTCTCCTTAAC<br>Reverse: CAGAAACTACAAAAGACAGGCAGTT |
| **Fmo2** | Mouse | Forward: GTCATTACCAACACTAGCAAAGAAATGT<br>Reverse: CTGAAACTGAATATATTTTAGGAGATCAAA |
| **Gpx3** | Mouse | Forward: AGTATGCAGGCAAATATATCCTCTTT<br>Reverse: AACATACTTGAGACTGGGGAGTATCT |
| **Hspa1a** | Mouse | Forward: TAAACTGTCTTTCAGTTACTTTGTGTATTG<br>Reverse: ACATATCTCTGTCTCTTTGTGTATTCTGAT |
| **Krt1** | Mouse | Forward: TATTAGAGAAAAGAGCTATGGTTAATTGC<br>Reverse: CTAGATCTGAAGTCCTCTTTGAAATGT |
| **Nos2** | Mouse | Forward: ATAGTTTCCAGAAGCAGAATGTGAC<br>Reverse: GTAGTAGAATGGAGATAGGACATAGTTCAA |
| **Nox4** | Mouse | Forward: CCAGAATACTACTACATTCACCAAATGTT<br>Reverse: CTGAGAAAATACAGATAGTTACACCACAT |
| **Noxa1** | Mouse | Forward: CTGCAGAGTATATGGAGGAATGTG<br>Reverse: TGTACTGAGCTACTACTTGGTAGAGGACT |
| **Nox2** | Mouse | Forward: AACTGTATGCTGATCCTGCTGC<br>Reverse: GTTCTCATTGTCACCGATGTCAG |
| **Kdm4a** | Mouse | Forward: TACTCTCTATGAACAGCACGTTGAT<br>Reverse: CTGTATAGATCCATGTCTTCCGTGT |
| **Kdm5c** | Mouse | Forward: GAGGAAAAGGACAAGGAATATAAACCC<br>Reverse: GTTCTGGATTCTTTTCAATGTCTTCCT |
| **Kdm6b** | Mouse | Forward: CCCTAACAAACCCTATTATGCTCCT<br>Reverse: CTCTGATTCATACAACTGTCCAAGC |
| **Dot1l** | Mouse | Forward: CTCCTGTGACAAGTGCATTTACTTT<br>Reverse: TGACTCTCTTATACACAGATGCCAG |
| **Ehmt2** | Mouse | Forward: GTTCATCTGCGAGTATGTAGGAGAG<br>Reverse: TAAACCTCGCCATCCTTGTTATCTAAAT |
| **Hdac4** | Mouse | Forward: GCTGAATCAAGTTGACATGAGGAAA<br>Reverse: CCAGAGCCATAGAAAATGGTTTTGT |
| **Hdac10** | Mouse | Forward: ATGGATTCTGTGTGTTCAACAATGT<br>Reverse: GGGTCATCGTTGAAGATATACTGGA |
| **Dnmt3a** | Mouse | Forward: AGCGTCACACAGAAGCATATCCAGGAG<br>Reverse: GGCCAGTACCCTCATAAAGTCCCTTGC |
| **18S** | Mouse | Forward: CGGCTACCACATCCAAGGAA<br>Reverse: GCTGGAATTACCGCGGCT |
| **MALAT1** | Human | Forward: CAACGAAGGCTTAAAGTAGGAC<br>Reverse: GCTGACACTTCTCTTGACCTTAG |
| **RPL4** | Human | Forward: GCCTGCTGTATTCAAGGCTC<br>Reverse: GGTTGGTGCAAACATTCGGC |
